# Modulation of Capsid Dynamics in Bromoviruses by the Host and Heterologous Viral Replicase

**DOI:** 10.1101/2022.08.16.504222

**Authors:** Antara Chakravarty, A. L. N. Rao

## Abstract

The tripartite genome of *Cowpea chlorotic mottle virus* is packaged into three morphologically indistinguishable icosahedral virions with T=3 symmetry. The two virion types, C1^V^ and C2^V^, package genomic RNAs 1 (C1) and 2 (C2), respectively. The third virion type, C3+4^V^, co-packages genomic RNA3 and its sub-genomic RNA (RNA4). In this study, each virion type was separately assembled *in planta* using a previously established agroinfiltration approach. The three virion types are indistinguishable under the EM and by electrophoretic mobility. However, the stability and the capsid dynamics evaluated respectively by the differential scanning fluorimetry and MALDI-TOF analysis following trypsin digestion distinguished the three virions type. Proteolytic analysis revealed that, while retaining the structural integrity, C1^V^ and C2^V^ virions released peptide regions encompassing the N-terminal arginine-rich RNA binding motif. In contrast, a minor population of the C3+4^V^ virion type was sensitive to trypsin-releasing peptides encompassing the entire capsid protein region. In addition, we also evaluated the effect of the two host organisms on the capsid dynamics. The wild-type CCMV virions purified from cowpea are highly susceptible to trypsin digestion, while those from *N. benthamiana* remained resistant. Finally, the MALDI-TOF evaluated the relative dynamics of C3+4^V^ and B3+4^V^ virions assembled under the control of the homologous vs. heterologous replicase. The significance of the viral replicase in modulating the capsid dynamics was evident by the differential sensitivity to protease exhibited by B3+4^V^ and C3+4^V^ virions assembled under the homologous vs. heterologous replicase. The significance of these results in relation to viral biology is discussed.

**IMPORTANCE:** Infectious virus particles or virions are considered static structures and undergo various conformational transitions to replicate and infect many eukaryotic cells. Although viral capsid fluctuations, referred to as dynamics or breathing, have been well studied in RNA viruses pathogenic to animals, such information is limited among plant viruses. *Cowpea chlorotic mottle virus* (CCMV) and Brome mosaic virus (BMV), RNA viruses pathogenic to plants, contain a segmented genome distributed among three physically and structurally homogenous icosahedral virion types. This study examines the role of the host organism and the heterologous viral replicase in modulating the thermal stability and capsid dynamics in CCMV and BMV. The results accentuate that alteration of the capsid dynamics by the host type and heterologous replicase are likely to affect the biology of the virus.

## INTRODUCTION

The assembly of viral capsids is promoted and directed by the interaction between the capsid protein (CP) subunits and the viral nucleic acid in both sequence-specific and non-specific manner with the assistance of other scaffolding proteins (1, 2). Once matured, the primary function of a viral capsid is to encapsidate the viral genome and remain stable enough to withstand the physicochemical environmental conditions until a susceptible host cell is encountered. In addition, assembled viral capsids must be flexible enough to fulfill multifunctional roles such as receptor binding, cellular entry, navigating the intracellular environment, and release of the viral genome. To meet these requirements, viral capsids are conformationally dynamic assemblies fluctuating from inside to outside of the particle (2). These fluctuations or the protein subunits’ dynamism manifest the breathing of the virus particles that position the biologically relevant surface peptides critical for their infectivity (3).

The three single-stranded, positive-sense genomic RNA molecules of the two closely related members of the family *Bromoviridae,* namely *Cowpea chlorotic mottle virus* (CCMV; C1, C2, and C3) and *Brome mosaic virus* (BMV; B1, B2, and B3) are encapsidated separately into three individual virions (4). The two largest monocistronic genomic RNAs of CCMV and BMV-RNA1 (C1, 3171 nt and B1, 3237 nt) and- RNA2 (C2, 2774 nt and B2, 2864 nt), encoding nonstructural replicase proteins 1a (methyltransferase/helicase-like) and 2a (RNA-dependent RNA polymerase) respectively, are packaged separately into two virion types, C1^V^/B1^V^ and C2^V^ /B2^V^ (4). The dicistronic genomic RNA3 (C3 and B3) encodes a nonstructural movement protein (MP) at its 5’ and a structural CP at its 3’ end. In these two viruses, CP is translated from a replication-derived subgenomic RNA4 (C4 and B4) via an internal initiation mechanism from the genomic progeny minus-strand RNA3. Genomic RNA3 and subgenomic RNA4 (sgRNA4) are co-packaged into a third virion type: C3+4^V^ and B3+4 (5).

BMV and CCMV exhibit similar genome organization and replication mechanisms (6, 7). However, they do differ in many properties, including the required virus-encoded genes for cell-to-cell spread (8), electrophoretic mobility patterns of respective virions (9), and finally, the interaction of highly conserved N-terminal basic arm region of the CP with respective genomic and subgenomic RNAs during encapsidation (10, 11). The CPs of CCMV and BMV are 190 and 189 amino acids in length (aa), respectively, and are 70% homologous. In these two viruses, the first N-terminal 25 aa region of the CP is not visible in the crystal structure since this highly basic region projects into the interior region interacting with the RNA, while the remainder of the CP is highly structured (4). The interaction between amino and carboxyl termini is essential for forming CP dimers, the building blocks for the virus assembly (12).

Fully assembled virions of CCMV and BMV exhibit icosahedral morphology with a diameter of ∼28 nm and T=3 quasi-symmetry (13, 14). Although the three genomic and a single subgenomic RNA are distributed disproportionately among three virions, they sediment as a single peek and are physically indistinguishable by any known procedures. Recently, we applied an *Agrobacterium*-mediated transient expression system (agroinfiltration) to independently assemble each of the three virion types of BMV (9). Application of limited proteolysis in conjunction with MALDI-TOF analysis revealed that B1^V^ and B2^V^ virions are distinct from the B3+4^V^ in their stability and dynamics, suggesting that RNA-dependent capsid dynamics play an important biological role in the viral life cycle.

Although at biochemical and structural levels, CCMV and BMV are similar, at biological and genetic levels, they exhibit the following two differential traits (i) Inherently, CCMV and BMV are respectively adapted to dicotyledonous Cowpea (*Vigna sinensis*) monocotyledonous plants (brome grass); (ii) hybrids viruses involving the exchange of the genomic RNA3, but not those encoding the replicase genes, are biologically viable (15). Keeping these two differential traits in perspective, in this study, we comparatively evaluated the following characteristics: (i) thermal stability and capsid dynamics of each virion type of CCMV and compared to those of BMV; (ii) the influence of the host organism on the dynamics of WT CCMV virions and finally (iii) thermal stability and capsid dynamics of virions of B3+4^V^ and C3+4^V^ assembled under the control of heterologous replicase. We observed a remarkable difference in the thermal stability and capsid dynamics within and among the three virion types of CCMV and BMV. In addition, we also observed that the type of host organism and heterologous replicase have a profound influence on the dynamics of the assembled virions.

## RESULTS

Throughout this study, individual virions packaging the CCMV genomic RNA1(C1), RNA2 (C2), and RNA3 (C3) plus subgenomic RNA4 (C4) are respectively referred to as C1^V^, C2^V^, and C3+4^V^. Likewise, those of BMV are designated as B1^V^, B2^V^, and B3+4^V^. Wild type (Wt) represents a mixture of three virion types of either CCMV or BMV, purified from infected plants.

### Characteristic properties of the three individually assembled virion types of CCMV (C1^V^, C2^V^, and C3+4^V^)

Fig.1 (Panel IA) summarizes the characteristic features of the transfer DNA (T-DNA)-based vectors engineered to express the three biologically active genomic RNAs (gRNAs) of CCMV (pC1, pC2, and pC3). Fig. 1 (Panel IB) summarizes the characteristic features of the plasmids engineered to transiently express CCMV replicase proteins p1a (pC1a), p2a (pC2a), and the CP (pCCP). Using these constructs, the strategy (Fig. 1, Panel II) used for assembling the three individual virion types C1^V^, C2^V^, and C3+4^V^, is essentially the same as that was described previously (9) for assembling BMV virion types using the constructs described in Fig. 1 (Panel IC, D). The following five sets of the inocula resulted in the assembly of the three virion types of CCMV *in planta* (i.e., C1^V^, C2^V^, and C3+4^V^), *<u>Set 1</u>*: pC1+pC2+pC3 to induce wild-type infection resulting in the accumulation of all three virion types that served as a positive control (Fig. 1, Panel IIA); *<u>Set 2</u>*: pCCP to transiently express CCMV CP only that served as a negative control (Fig. 1, Panel IIB); *<u>Set 3</u>*: pC1+pC2a+pCCP to assemble virion type C1^V^ (Fig. 1, Panel IIC); *<u>Set</u> <u>4</u>:* pC1a+pC2+pCCP to assemble virion type C2^V^ (Fig. 1, Panel IID) and *<u>Set</u>* <u>5</u>: pC1a+pC2a+pC3 to assemble virion type C3+4^V^ (Fig. 1, Panel IIE).

**FIG 1.**
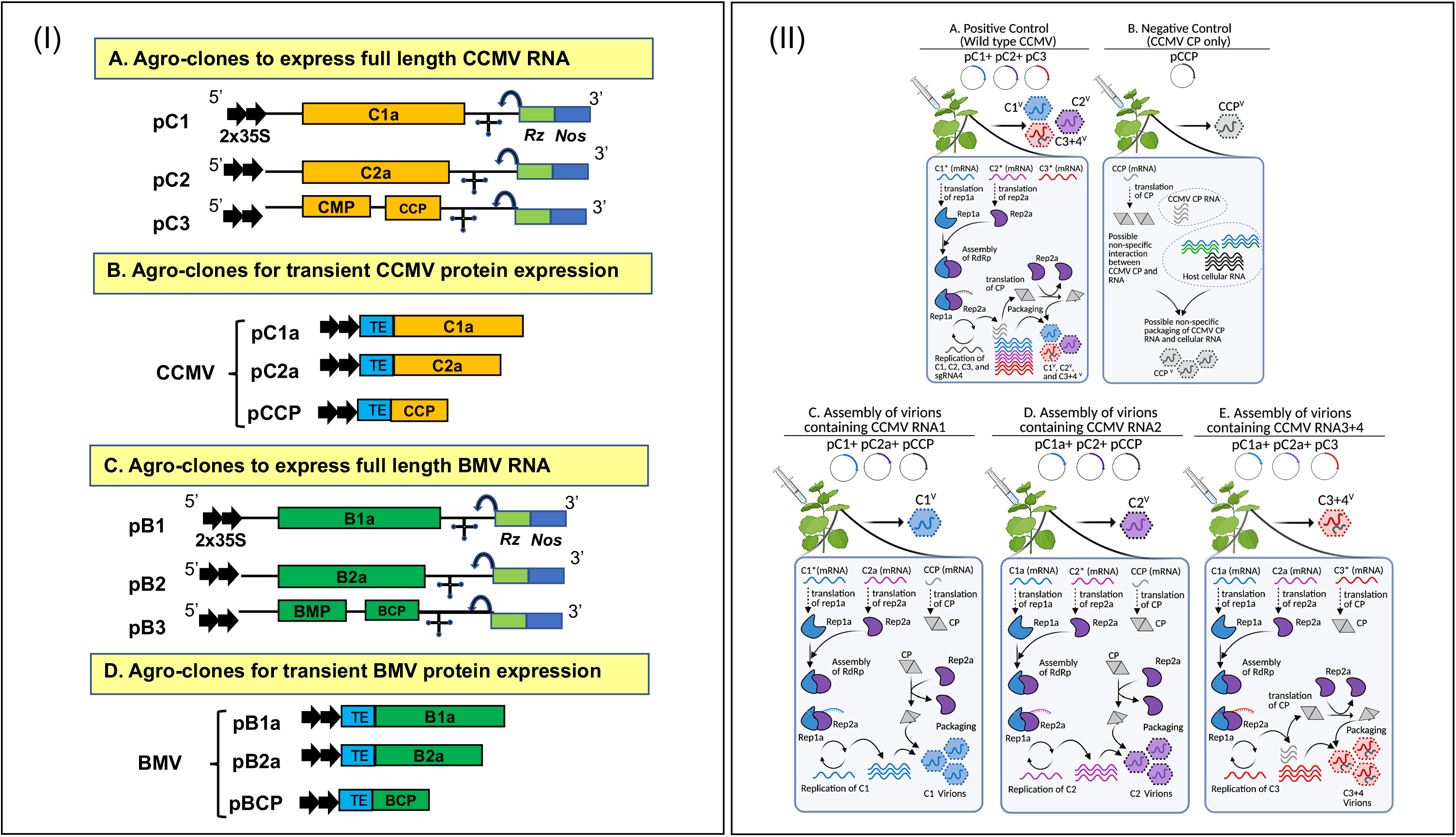
Agroplasmids and agroinfiltration *in planta*. *Panel I*. (A) Characteristics of T-DNA-based agroplasmids for the transient expression of full-length CCMV genomic RNAs in plants as described previously for BMV (9). The 35S-C1 (pC1), 35S-C2 (pC2), and 35S-C3 (pC3) constructs respectively contain the full-length cDNA copies of CCMV genomic RNA1 (C1), -2 (C2), and -3 (C3). Single lines and solid boxes represent noncoding and coding regions, respectively. Monocistronic C1 encoding replicase protein 1a (C1a) and monocistronic C2 encoding replicase protein 2a (C2a) are indicated. In C3, the locations of the movement protein (CMP) and capsid protein (CCP) genes are shown. At the 3’ end of each construct, the clover leaf-like structure represents a tRNA-like motif conserved among all three CCMV genomic RNAs. A set of two black arrows at the 5’ ends represent the double 35-S promoter. A bent arrow indicates the predicted self-cleavage site by the ribozyme. The location of the *Nos* terminator is indicated. (B) Agroconstructs for transient expression of C1a, C2a, and CP (pCCP). Open reading frames (ORFs) of CCMV p1a, p2a, and CP were fused in-frame to a binary vector using *Stu*I and *Spe*I sites. (C) Characteristics of agroplasmids for the transient expression of full-length BMV genomic RNAs in plants as described previously (9). (D) Features agroconstructs for the transient expression of BMV replicase proteins (B1a and B2a), and CP (pBCP) as described previously (9). *Panel II*. Agroinfiltration strategy used for the assembly of each virion type of CCMV. (A) (Positive control) Infiltration of the inoculum containing a mixture of pC1+pC2+pC3 would result in the induction of Wt CCMV infection containing a mixture of all three virions (i.e., C1^V^, C2^V^, C3+4^V^). (B) (Negative control) Infiltration of the inoculum containing agroplasmid pCCP would result in the expression of CP mRNA for the translation of Wt CP and its non-specific assembly of virions containing CP mRNA and cellular RNA. (C) Assembly of virions packaging CCMV RNA1 (C1^V^). A mixture of inoculum containing agrotransformants pC1+pC2a+pCCP (see Fig. 1 panel I for details) is infiltrated into *N. benthamiana* leaves to result the following. (i) Transcription of pC1 results in the synthesis of a biologically active full-length genomic RNA1 for the translation of functional replicase protein 1a; (ii) agrotransformant pC2a results in an mRNA competent for the translation of replicase 2a but not replication because it lacks the 5’ and 3’ noncoding regions and finally (iii) translation of the mRNA transcribed from agrotransformant pCCP gives the CP subunits for directing virion assembly. Assembly of functional replicase results from the interaction of proteins 1a and 2a catalyzes the replication of C1 RNA, followed by its packaging into virions by the transiently expressed CCMV CP subunits. (D) Assembly of virions packaging CCMV RNA2 (C2^V^). The inoculum shown in this panel is identical to the one shown in panel C, except that pC1 and pC2a are replaced by pC1a and pC2, respectively. Agroinfiltration of this inoculum results in the assembly of virions containing C2 RNA. (E) Assembly of virions packaging CCMV RNA3 and -4 (C3+4^V^). The inoculum shown in this panel is formulated to assemble virions packaging C3 RNA and sgC4 by the infiltration of a mixture of pC1a, pC2a, and pC3 agrotransformants. Transiently expressed replicase proteins 1a and 2a directs the replication of C3, followed by the synthesis of sgC4 for CP production. Fig1 panel II was created with BioRender.com.

Fig. 2 summarizes the progeny analysis from the five sets of inocula as mentioned above. Western blot analysis (Fig. 2A) confirmed the efficient expression of the monomeric and dimeric forms of the CCMV CP with expected size in plants infiltrated with each set (Fig. 2A, lanes 1-5). Next, virions purified from infiltrated leaves were analyzed for their morphology and size by the EM and their relative electrophoretic mobility pattern by agarose gel-electrophoresis (Fig. 2B, C). EM analysis (Fig. 2B) revealed that virion preparations of C1^V^, C2^V,^ and C3+4^V^ are indistinguishable from those of Wt CCMV isolated from cowpea or *N. benthamiana* in morphology and size (∼28-30 nm). Finally, when electrophoresed in agarose gels, samples of C1^V^, C2^V^, and C3+4^V^ virion types migrated as a single band (Fig. 2C). They were identical to that of Wt CCMV (migrating toward positive polarity) and distinct from that of control virions of BMV (migrating toward negative polarity). In addition, we also confirmed the nature of the packaged RNA in C1^V^, C2^V^, and C3+4^V^ by RT-PCR (data not shown). Furthermore, as expected, an inoculum containing all three virion types (i.e., C1^V^+ C2^V^ + C3+4^V^) remained infectious to plants. None of the pair-wise combinations (i.e., C1^V^+ C2^V^ or C1^V^+ C3+4^V^ or C2^V^+ C3+4^V^) remained non-infectious (data not shown). These results confirmed the purity and the authenticity of the three virion types of CCMV assembled independently in *N. benthamiana* plants.

**FIG 2.**
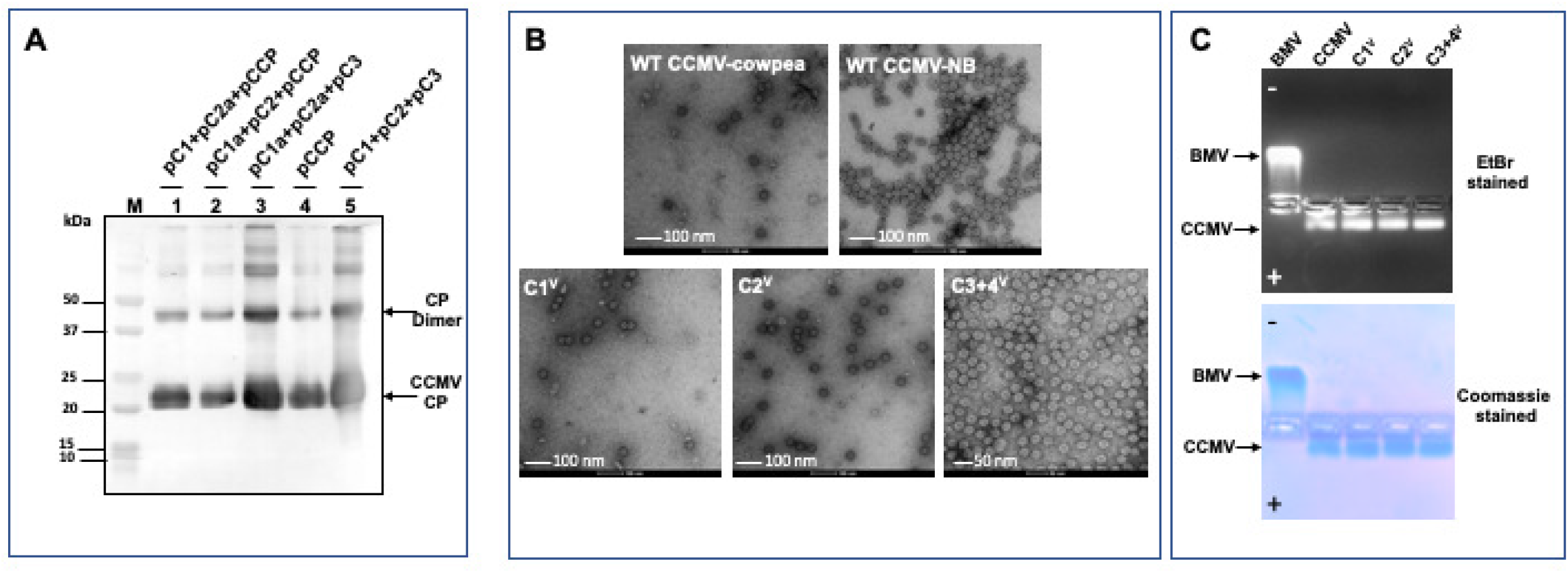
Progeny analysis of the three individually assembled virion types of CCMV (C1^V^, C2^V^, and C3+4^V^). *(A) Western blot analysis*. Total protein preparations from *N. benthamiana* leaves at 4 dpi were extracted and analyzed using an anti-CCMV CP antibody*. Lane 1*, pC1+pC2a+pCCP resulting in the assembly of C1^V^; *Lane 2,* pC1a+pC2+pCCP resulting in the assembly of C2^V^; *Lane 3,* pC1a+pC2a+pC3 resulting in the assembly of C3+4^V^; *Lane 4*, pCCP resulting in the assembly CCP^V^; *Lane 5*, pC1+pC2+pC3 resulting in the assembly of Wt CCMV infection (i.e., a mixture of C1^V^, C2^V^, and C3+4^V^). In each lane, a sample containing 30 μg of total leaf protein (estimated by the Bradford protein assay) was loaded. *(B) EM analysis*. Electron micrographs of the density gradient-purified virions of Wt CCMV from either *N. benthamiana* or cowpea plants or autonomously assembled virions of C1^V^, C2^V^, and C3+4^V^ via agroinfiltration. Virion samples were negatively stained with uranyl acetate prior to examining under the EM. The scale bar is indicated on each image. *(C) Virion electrophoresis*. Density gradient-purified virions of Wt BMV (lane 1), Wt CCMV (lane 2), C1^V^ (lane 3), C2^V^ (lane 4), and C3+4^V^ (lane 5) were subjected to electrophoresis in an agarose gel as described under Materials and Methods. The gel shown on the top panel was stained with ethidium bromide to detect RNA and then re-stained with Coomassie Brilliant Blue R-250 to detect the protein (bottom panel). The positions of BMV and CCMV virions migrating toward negative and positive, respectively, are indicated.

### Stability differences among C1^V^, C2^V^, and C3+4^V^ virions

We previously observed that differential scanning fluorimetry (DSF) is ideal for comparing the relative stability of the three virion types in BMV (9). Fig. 3 (A-D) summarizes the results of the DSF analysis of the temperature-dependent melting for C1^V^, C2^V^, and C3+4^V^ and control samples of lysozyme under two buffer and pH conditions. First, we compared the temperature-dependent melting of C1^V^, C2^V^, and C3+4^V^. Although each virion type revealed near identical thermal melting profiles displaying a single unstable peak at ∼70°C in the virus suspension buffer (pH 4.5) (Fig. 3A), a clear distinction in the thermal instability among the three virion types was evident in the phosphate buffer (pH 7.2). For example, C1^V^ and C2^V^ displayed two major thermally stable population peaks (Fig. 3B; 46°C and 88°C for C1^V^ and 50°C and 88°C for C2^V^). By contrast, for C3+4^V^, 80% virion population showed instability at 48°C while the remaining 20% melted at 82°C (Fig. 3B, compare two peaks of C1^V^ and C2^V^ vs. C3+4V). As expected, the melting profile of the lysozyme control (Fig. 3 C, D) remained indistinguishable either in the virus suspension buffer (pH 4.5) or phosphate buffer (pH 7.2). Interestingly, these melting profiles of C1^V^, C2^V^, and C3+4^V^ are distinct from those of BMV (see Discussion).

**FIG 3.**
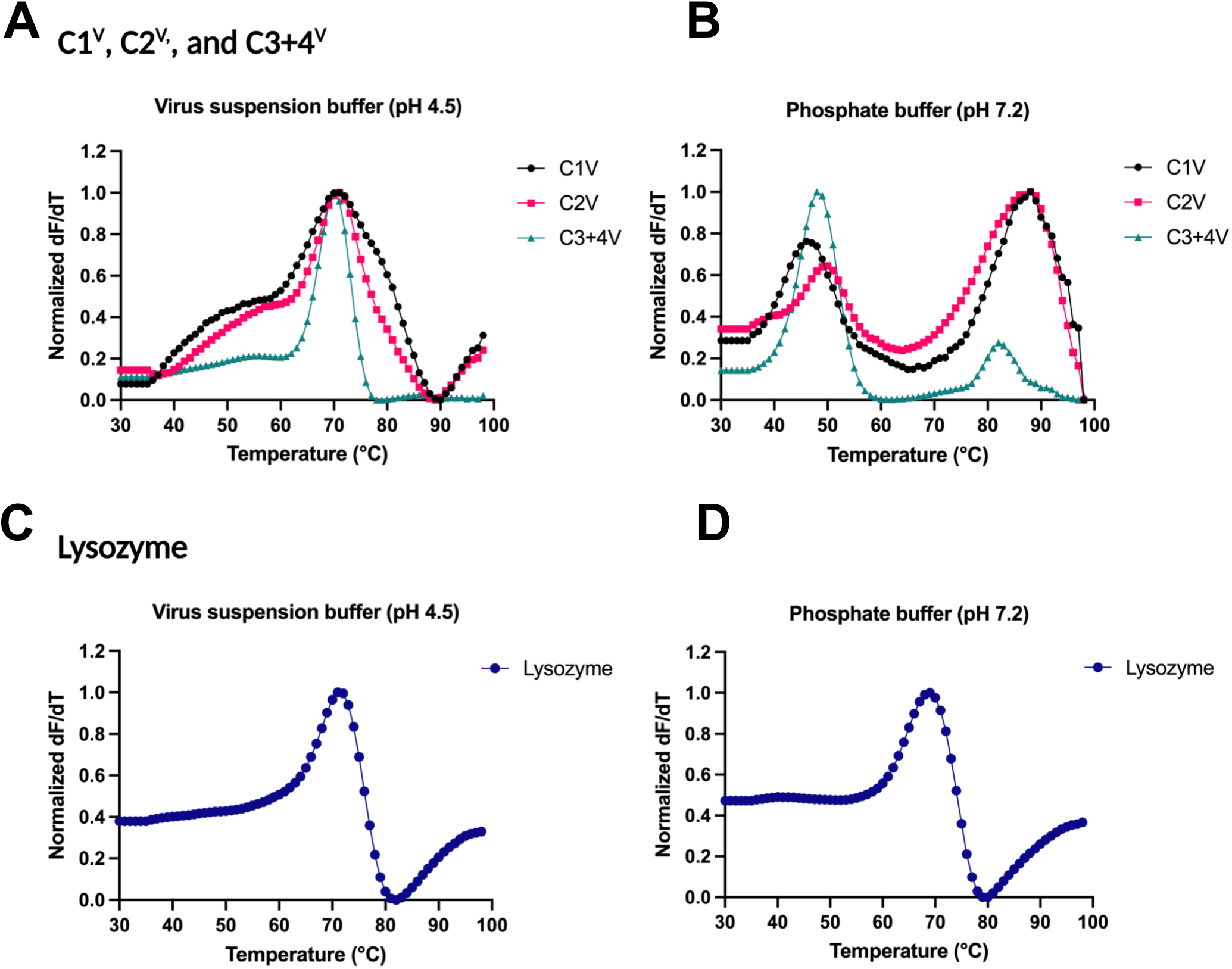
Stability analysis of the three CCMV virion types using differential scanning fluorimetry (DSF). DSF measurements of derivative of fluorescence intensity as a function of temperature for each virion type normalized so that the maximum of the derivative is set to 1 (y-axis) during heating of (A and B) C1^V^, C2^V^, and C3+4^V^ and (C and D) lysozyme (control) at various temperatures (x-axis) under indicated conditions. Panel A: Virions of C1^V^, C2^V^, and C3+4^V^ displayed identical temperature dependence (∼70°C) for melting under virus suspension buffer conditions at pH 4.5. Panel B: At pH 7.2, a portion of virions of C1^V^, C2^V^, and C3+4^V^ melted at lower temperatures (∼50°C), whereas C1^v^ and C2^v^ displayed another major thermally unstable population of peaks at ∼90°C (see text for details). Panel C and D: Lysozyme used as positive control showed identical melting characteristics (∼70°C) under both pH conditions. The figures indicate representative data from N=3 independent experiments.

### Capsid dynamics of the three virion types of CCMV

As observed in BMV (9), in CCMV also, the three virion types (C1^V^, C2^V^, and C3+4^V^) are accumulated disproportionately (46% C1^V^+C2^V^ vs. 54% C3+4^V^; data not shown). To verify whether physically indistinguishable three virion types of CCMV are dynamically distinct, we evaluated the capsid dynamics of each virion type by MALDI-TOF analysis following trypsin digestion (16). Virion preparations of C1^V^, C2^V^, C3+4^V^, and CCPV assembled in *N. benthamiana* were concurrently subjected to trypsin digestion for 10, 30, 45 min, and 3 hr. Then, each sample was divided into three equal aliquots and used for Western blot, EM and MALDI-TOF analyses. Western blot identified the cleavage peptides, EM analysis verified the structural integrity and MALDI-TOF identified the cleavage sites. Undigested virion samples served as controls.

Fig. 4 (A, B) summarizes these results. Western blot data shown in Fig. 4A revealed that undigested samples of the four individual virion types (i.e., C1^V^, C2^V^, C3+4^V,^ and CCPV) migrated as a single intact band with the expected molecular weight (∼20 kDa). Analysis of the trypsin cleavage products for each virus sample revealed exciting profiles. Trypsin digestion of the three of four virion types, C1^V^ and C2^V^, and CCP^V^ for 10, 30, and 45 min are indistinguishable from the undigested control samples (Fig. 4A; Lanes 5, 6, 8 for 10 min; lanes 9, 10, 12 for 30 min and lanes 13, 14, and 16 for 45 min). By contrast, for the C3+4^V^ virion type, although a majority (>50%) of the protein remained intact, several faster-migrating peptide fragments were consistently detected (Fig. 4A, lanes 7, 11, and 15). Extending the trypsin digestion time to 3h did not change these profiles (Table 1). EM analysis of the trypsin-digested virion preparations (Fig. 4B) confirmed the Western blot data, showing that most of the C1^V^ and C2^V^ (Fig, 4B), as well as CCP (data not shown), remained structurally intact. Whereas for C3+4^V^ virions, the majority of the virions remained intact, while structurally disrupted virions are also visible (Fig, 4B).

**FIG 4.**
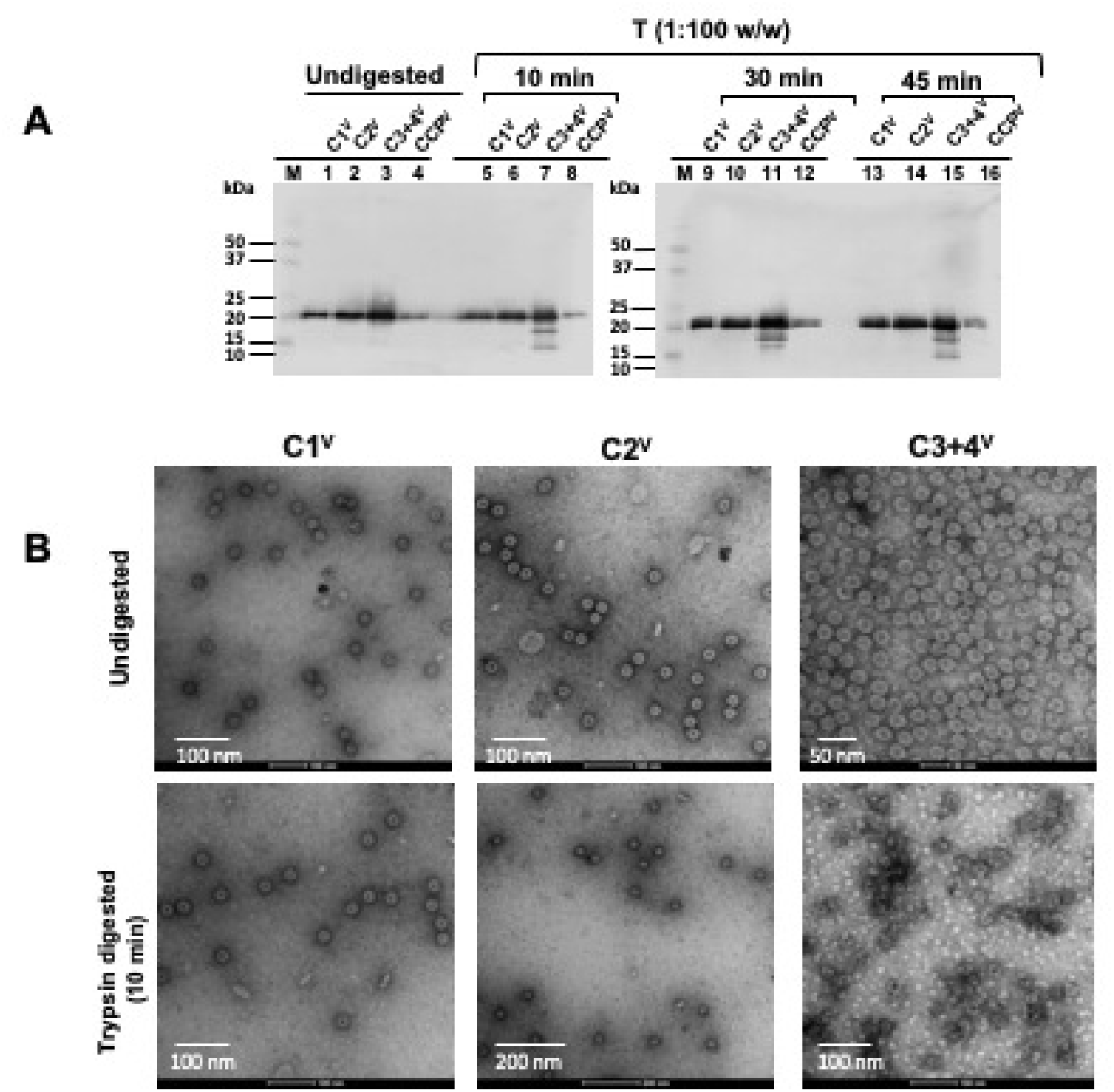
Proteolysis of C1^V^, C2^V^, C3+4^V^, CCP^V^. *(A) Western blot analysis*: Virion preparations of either C1^V^ or C2^V^ or C3+4^V^ or CCP^V^ were either undigested or digested with trypsin for 10 min, 30 min, or 45 min and subjected to Western blot analysis as described under the Materials and Methods section. *(B) EM analysis:* Negative-stain electron micrographs showing the integrity of the undigested and trypsin-digested virion preparations of the indicated samples. The scale bar is indicated on each image.

**TABLE 1.**
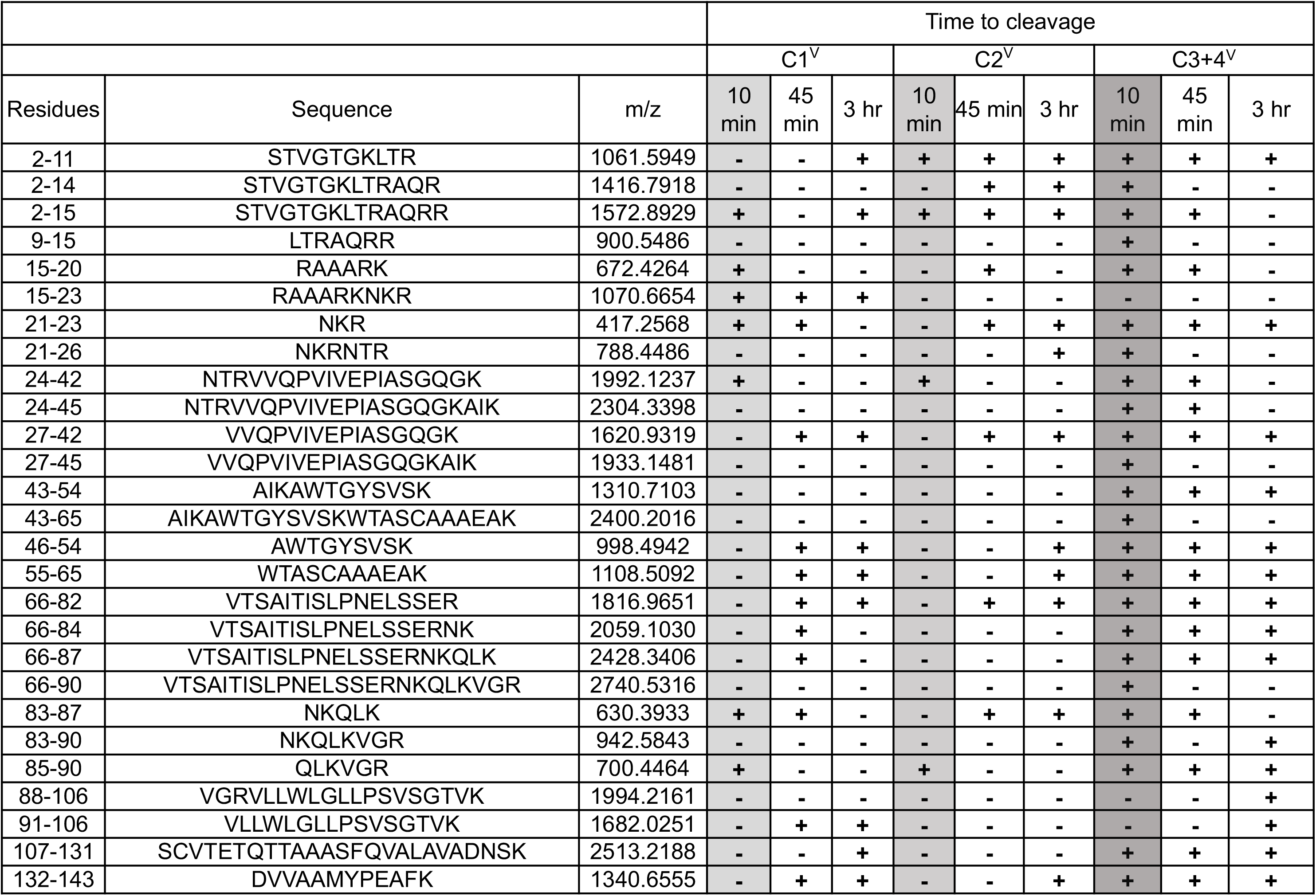
Kinetics of trypsin cleavage sites located on C1^V^, C2^V^, and C3+4^V^. Amino acid fragments released during the early time points by C1^V^, C2^V^ are highlighted in light grey, and those released by C3+4^V^ are highlighted in dark grey.

Fig. 5 (A, B) summarizes the organization of the three CP subunits (A, B, and C) on the surface of the CCMV virion and T=3 lattice. Fig. 5 (C-E) summarizes the position of the trypsin cleavage sites located on each CP subunit. The deconvoluted mass spectrum obtained from the total ion chromatogram (TIC) of C1^V^, C2^V^, and C3+4^V^ indicated that all the virions are assembled from a single protein of 20 kDa (data not shown). Analysis of the peptides released by the trypsin digestion at early (10 min) and late time points (45 min and 3 hr) distinguished virions of C1^V^ and C2^V^ from those of C3+4^V^ (Fig 5 F-H). For example, 10 min after the digestion, C1^V^ and C2^V^ virions appeared to release a small proportion of the peptides corresponding to aa encompassing the N-ARM region (e.g., aa 2-15; Fig. 5F; Table 1). Since the N-proximal region is predicted to be mobile/disordered and internalized following its interaction with RNA (17) and are not ordered in the crystal structure, their cleavage did not affect the structural integrity of most of the C1^V^ and C2^V^ virions, as seen under EM (Fig. 4B). Also, there were peaks of very low-intensity corresponding to aa 83-87 and 85-90 (Table 1). These fragments could have been generated from a very small percentage of virions, hence not detected in Western blot. The trypsin digestion to 45 min and 3 hr of C1^V^ and C2^V^ released several other aa fragments corresponding to additional cleavage sites on the virion surface (Table 1), including K65 (Fig. 5 C-E). These fragments are present at a low % intensity and hence are not detectable by the Western blot (Fig. 4A).

**FIG 5.**
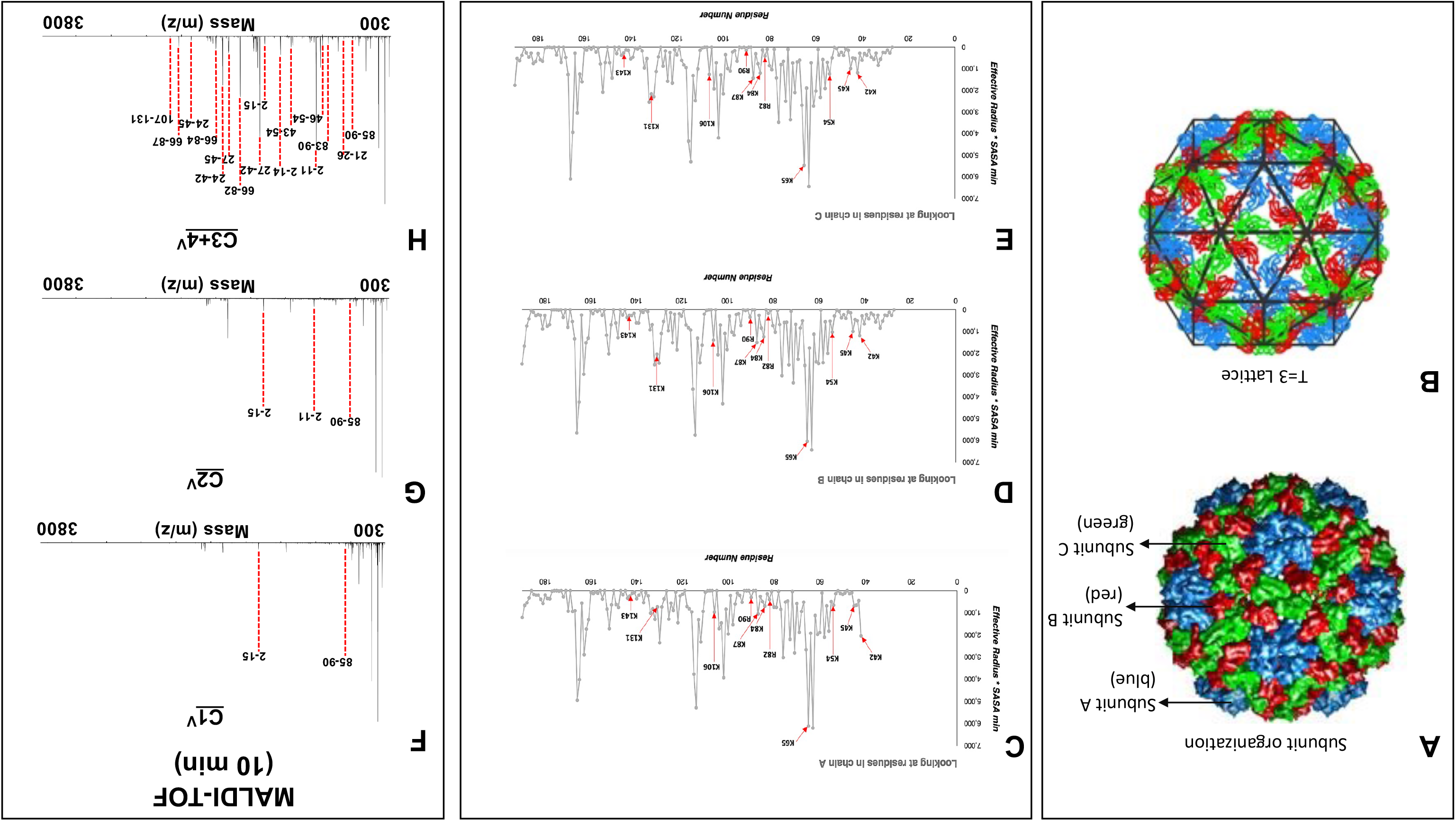
The organization of the three CP subunits (A, B, and C) on the surface of the CCMV virions and analysis of the peptides released by the trypsin digestion of virions. (A and B): Structure of CCMV capsid based on the PDB entry 1CWP showing the A, B, C subunits and T=3 lattice. (C-E): The position of the trypsin cleavage sites located on each CP subunit A, B, and C. (F-H) MALDI-TOF analysis of peptides released from virions of either C1^V^ or C2^V^ or C3+4^V^ following digestion with trypsin at the indicated time point. Peaks are labeled with corresponding polypeptide fragments of predicted amino acid residues. Table 1 summarizes masses and identifies the corresponding amino acid residues.

In contrast, irrespective of the time of incubation, virions of C3+4^V^ remained susceptible to trypsin digestion. As early as 10 min, virions of C3+4^V^ released multiple peptides encompassing the entire CP region (Fig. 5 H). This digestion pattern did not change significantly at later time points (45 min and 3 hr). Mass mapping identification of the released peptides revealed that the accessible trypsin cleavage sites (e.g., K45, K54, K65, K87, K106, and K131; Fig. 5 C-E) (Table 1) were consistent with the reported surface structure of the CCMV virions (http://viperdb.scripps.edu/). Given the importance of the C-terminal peptide region in the virus assembly (16), these observations explain why some of the virions of C3+4^V^ appeared degraded under EM (Fig. 4B). Taken together, it is reasonable to conclude that a percentage of trypsin-digested C3+4^V^ samples show qualitatively distinct capsid dynamics from those of C1^V^ and C2^V^, exhibiting a different conformational arrangement of the CP amino acid residues in the virion exterior.

### Effect of the host type on the stability of CCMV virions

For CCMV, cowpea is the natural host, while *N. benthamiana* is a symptomless experimental host (18). Since the type of host plant can have a profound influence on virion phenotype (19), we wanted to examine the effect of the host plant on the stability and dynamics of CCMV virions. Consequently, we compared the stability of CCMV virions purified from cowpea and *N. benthamiana* plants using DSF and dynamics by MALDI-TOF following trypsin digestion. For the first assay, CCMV virions purified from cowpea and *N. benthamiana* were suspended in either virus suspension buffer (pH 4.5) or phosphate buffer (pH 7.2) and subjected to DSF analysis. Fig. 6A summarizes these results. In virus suspension buffer (pH 4.5), most of the Wt CCMV virion population purified from cowpea and *N. benthamiana* plants displayed a single melting peak at 65°C and 69°C, respectively. In addition, virions purified from *N. benthamiana* but not from cowpea plants (Fig. 6A, left panel) revealed a small melting population appearing as a shoulder peak at ∼50°C. By contrast, DSF analysis in the phosphate buffer (pH 7.2) revealed a contrasting scenario (Fig. 6A, right panel). For example, unlike virus suspension buffer (pH 4.5), CCMV virions from cowpea plants displayed a significant peak at 46°C and a minor peak at 87°C (Fig. 6A, right panel). In contrast, CCMV virions obtained from *N. benthamiana* plants displayed two significant peaks with equal intensity, one at 45°C and the other at 79°C (Fig. 6A, right panel).

**FIG 6.**
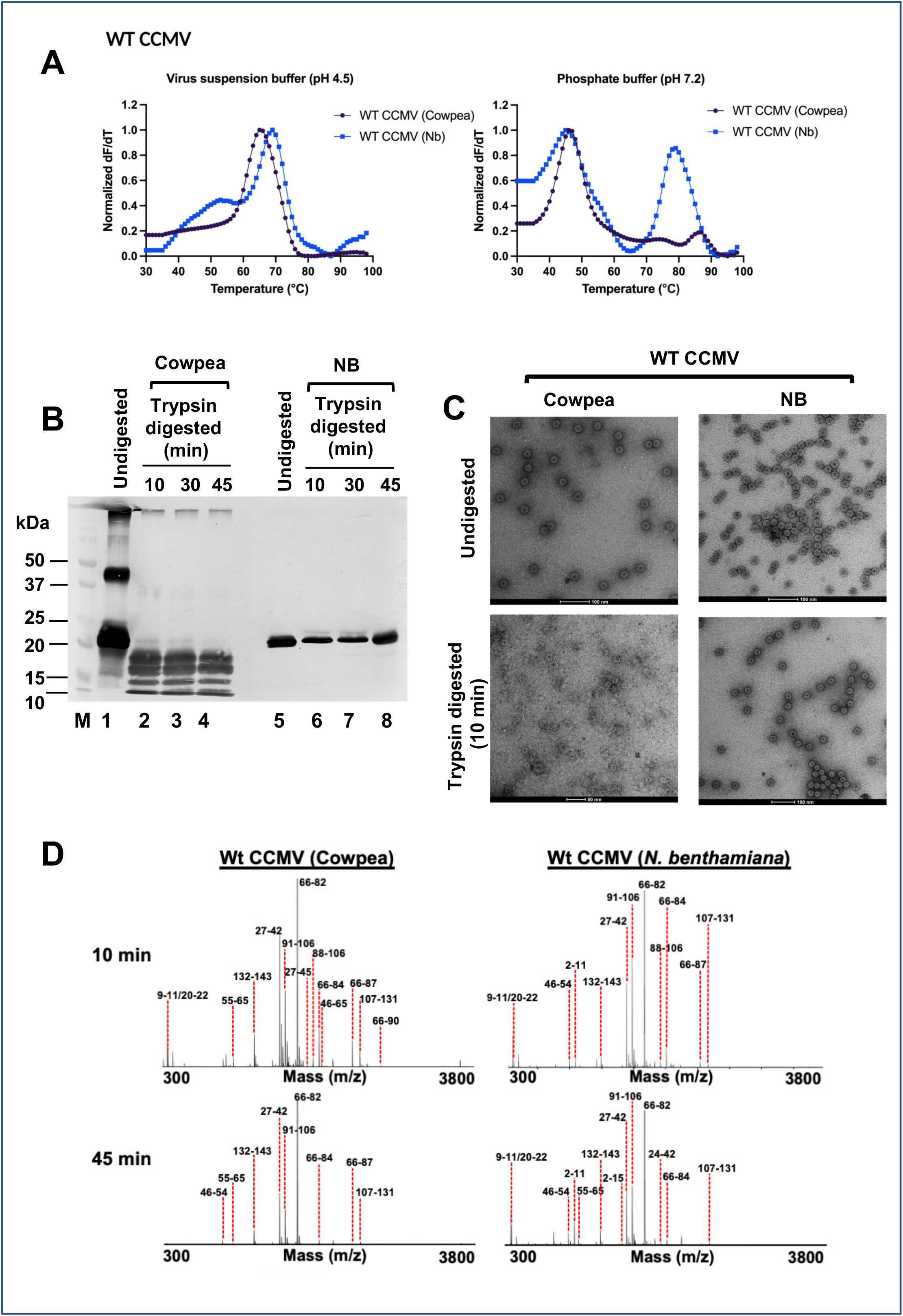
Effect of the host on the stability of Wt CCMV virions. (A) Stability analysis of the Wt CCMV virions isolated from either cowpea or *N. benthamiana* leaves using differential scanning fluorimetry (DSF). (Left panel) Most of the Wt CCMV virions purified from both host plants displayed a single peak melting at ∼70°C at pH 4.5. (Right panel) At pH 7.2, CCMV virions purified from cowpea displayed a major peak at ∼50°C and a minor peak at ∼90°C, whereas CCMV virions obtained from *N. benthamiana* plants displayed two major peaks with equal intensity, one at ∼50°C and the other at ∼80°C. (B) Western blot analysis of Wt CCMV from cowpea and *N. benthamiana* (NB). Prior to Western blot analysis, each virion preparation was either undigested or digested with trypsin for 10 min, 30 min, or 45 min. (C) Negative-stain electron micrographs show the integrity of the undigested and trypsin-digested virion preparations of Wt CCMV from cowpea and *N. benthamiana* (NB). (D) MALDI-TOF analysis. Wt CCMV virions from cowpea and *N. benthamiana* are subjected to MALDI-TOF analysis following trypsin digestion for 10 min as described in the Methods section. Peaks are labeled with corresponding polypeptide fragments of predicted amino acid residues.

Next, we evaluated the relative susceptibility of Wt CCMV virions from cowpea and *N. benthamiana* to trypsin digestion as described in the Methods section. Undigested virion preparation served as control, and Fig. 6B summarizes these results. When compared to the undigested control sample (Fig. 6B, lanes 1 and 5), Wt CCMV virions from cowpea plants are susceptible to trypsin digestion as early as 10 min (Fig. 6B, lanes 2-4), while those from *N. benthamiana* plants remained stable even after 45 min (Fig. 6B, lanes 6-8). EM analysis (Fig. 6C) confirmed these observations. To identify the peptide fragments, MALDI-TOF analysis was performed for Wt CCMV virion from cowpea and *N. benthamiana*. Results are shown in Fig. 6D. As expected, Wt CCMV virions from cowpea released peptide fragments encompassing the entire CP region (Fig. 6D, left panel). Although Western blot analysis failed to detect any cleavage products when Wt CCMV virions from *N. benthamiana* were subjected to trypsin treatment, MALDI-TOF revealed a profile similar to that of CCMV virions from cowpea plants. However, these peptides appeared to be weaker in intensity and hence are not detected by the Western blot analysis. No change in the MALDI-TOF profile was observed with prolonged incubation with trypsin (data not shown). These results suggest, CCMV virions in *N. benthamiana* accumulate as two virion populations: a minor population susceptible to trypsin and a major trypsin resistant population. These results collectively accentuate that the host plant type has a profound influence on the stability and dynamics of CCMV virions. The discussion section addresses the significance of these observations.

### Influence of viral replicase on the stability and capsid dynamics

Analysis of trypsin digestion profiles of BMV (9) and CCMV (Fig. 5; Table 1) revealed a minor but clear difference in the capsid dynamics of the respective viruses. Despite variation in the genome sequence (15), construction of the stable hybrid viruses between BMV and CCMV involving the exchange of genomic RNA3 but not RNAs 1 and 2 have been reported (15). *N. benthamiana* leaves were infiltrated with the desired mixture of the agro-inocula resulting in the assembly of two sets of hybrid viruses (Fig. 7A) to verify the effect of the heterologous replicase on the dynamics of the assembled hybrid virions. The first set of hybrid viruses involved the following combination of the inocula: pB1+pB2+pC3 or pC1+pC2+pB3. These inocula would result in the assembly of virions using either CCMV CP (i.e., CCMV virions packaging BMV RNAs 1 and 2 and CCMV RNA 3+4) or BMV CP (BMV virions packaging CCMV RNAs 1 and 2 and BMV RNA 3+4).

**FIG 7.**
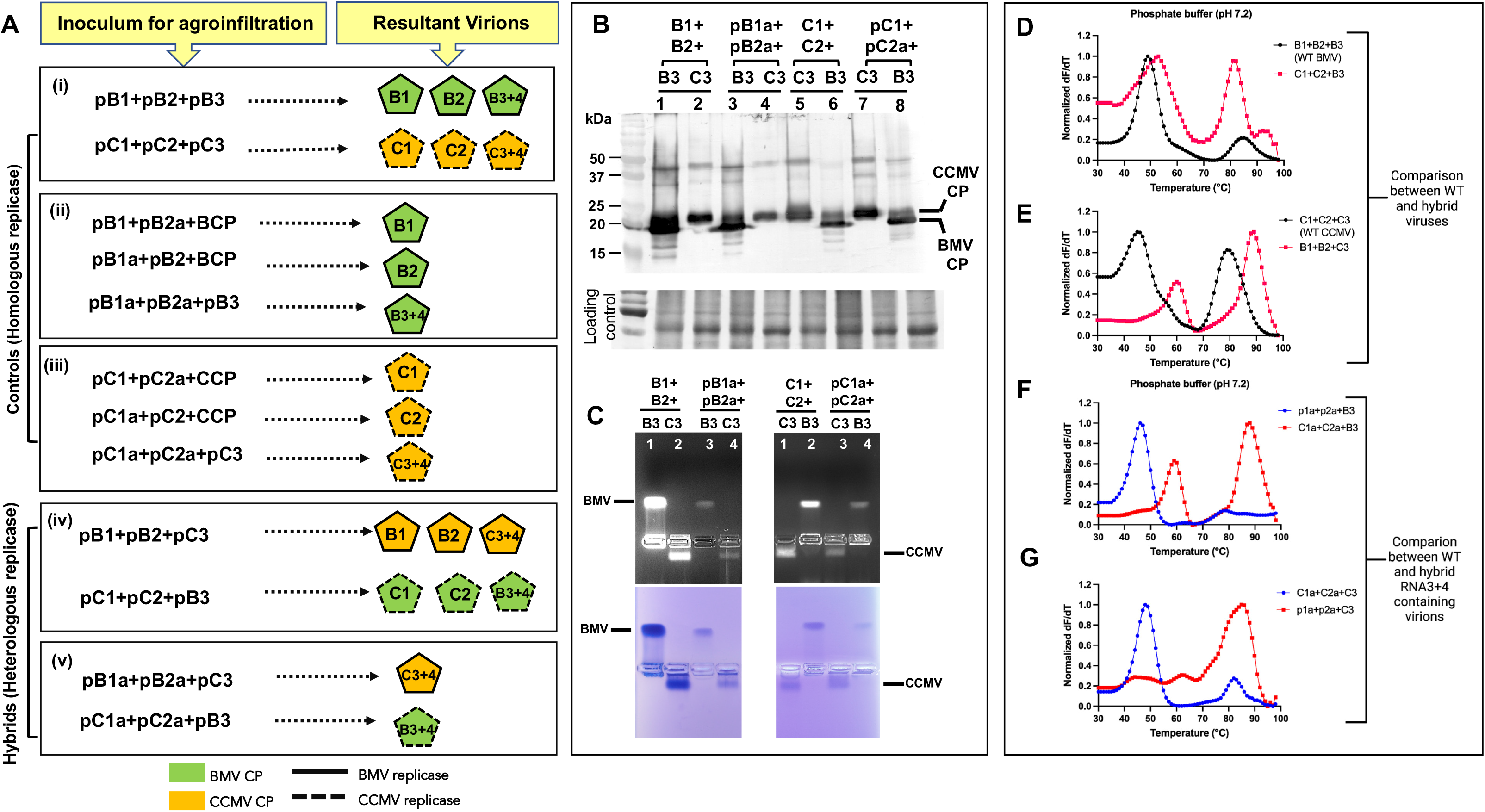
Properties of Wild-type and hybrid viruses constructed between BMV and CCMV. (A) Schematic representation of the strategy used for the (i) assembly of control virions of BMV Wt (pB1+pB2+pB3) and CCMV Wt (pC1+pC2+pC3); (ii) autonomous assembly of either B1^V^ or B2^V^ or B3+4^V^ as described previously; (iii) autonomous assembly of either C1^V^, C2^V^, and C3+4; (iv) hybrid virions assembled by the CCMV CP with BMV heterologous replicase: pB1+pB2+pC3 and hybrid virions assembled by the BMV CP with CCMV replicase: pC1+pC2+pC3; (v) assembly of virion type C3+4 with BMV replicase (pB1a+pB2a+pC3) and assembly of virion type B3+4 with CCMV replicase (pC1a+pC2a+pB3). (B) (Top panel) Western blot analysis. Lanes 1-8: Total protein preparations extracted from *N. benthamiana* at 4 dpi infiltrated with the indicated mixture of agrotransformants were subjected to Western blot analysis using anti-CP antibody against CCMV and BMV. Each lane has 30 μg of total protein estimated by Bradford protein assay. (Bottom panel) The gel is stained with Coomassie Brilliant Blue R-250 to demonstrate the loading controls. The samples are loaded in the order as mentioned above in lanes 1-8. (C) Electrophoretic mobility profiles: The following virion preparations were subjected to electrophoresis as described in the Materials and Methods section. Lanes 1 (left panel) Wt BMV or CCMV (right panel); Lanes 2 (left panel) hybrid virions of BMV assembled with CCMV CP, or hybrid virions of CCMV assembled with BMV CP (right panel); Lanes 3 left and right panel, respectively represent the virion type B3+4^V^ and C3+4^V^ assembled with homologous replicase; Lanes 4 left and right panel, respectively represent the virion type B3+4^V^ and C3+4^V^. The gels shown on the top panel were stained with ethidium bromide to detect RNA and then re-stained with Coomassie Brilliant Blue R-250 to detect protein (bottom panel). BMV and CCMV virions migrating toward negative and positive, respectively, are indicated. (D-G) DSF analysis: Stability analysis of the BMV and CCMV virions assembled in the presence of either homologous or heterologous replicase using differential scanning fluorimetry (DSF). DSF measurements of derivative of fluorescence intensity as a function of temperature for each virion type normalized so that the maximum of the derivative is set to 1 (y-axis) during heating of the virions indicated at various temperatures (x-axis) at pH 7.2. The virions assembled in the presence of either full-length RNAs or ORFs encoding heterologous replicase show unusual and increased thermal stability compared to those assembled as a result of interaction between homologous replicase and CP (see the text for details).

Genomic RNA3 is amenable for exchanging between BMV and CCMV (15). Therefore, we considered the assembly hybrid virions of RNAs 3+4 under heterologous replicase. Consequently, infiltration of pB1a+pB2a+pC3 would result in the assembly of C3+4^V^ by the CCMV CP generated by the BMV replicase. Likewise, infiltration of pC1a+pC2a+pB3 would result in the assembly of B3+4^V^ by the BMV CP generated by the CCMV replicase (Fig. 7A). Infiltration of the inocula resulting in the assembly of respective Wt viruses (BMV and CCMV) or a virion type packaging RNA 3+4 using homologous replicase (Fig. 7A; e.g., pB1+pB2+pB3; pC1+pC2+pC3 and pB1a+pB2a+pB3 and pC1a+pC2a+pC3) served as controls. Western blot analysis and electrophoretic mobility pattern (Fig. 7 B, C) confirmed the hybrid nature of the purified virions assembled under the direction of homologous and heterologous replicases.

The DSF analysis (Fig. 7 D, E) of the temperature-dependent melting of the two hybrid viruses and control samples (Wt BMV and Wt CCMV) was carried out under an environment that closely mimics the pH of living cells (phosphate buffer, pH 7.2). Virions of BMV and CCMV assembled in the presence of homologous replicase (i.e., B1+B2+B3 and C1+C2+C3) showed distinct thermal denaturation profiles. For example, most BMV virions melted at 49°C while a small population at 85°C (Fig. 7D; black peaks). By contrast, CCMV virions displayed two nearly equal populations: one melting at 45°C and the other at 79°C (Fig. 7E; black peaks). The role of viral replicase on the thermal profiles is evident when each hybrid virus was subjected to DSF analysis. Assembly of virions by the BMV CP synthesized by the heterologous replicase (C1+C2+B3; Fig. 7D) mimicked the melting profiles of Wt CCMV (C1+C2+C3; Fig. 7E). However, virions assembled with CCMV CP generated by the heterologous replicase (B1+B2+C3; Fig. 7E) appeared different in their thermal profiles compared to that of Wt BMV (B1+B2+B3; Fig. 7D). This variation perhaps can be attributed to the disproportionate accumulation of all three virion types.

The effect of heterologous replicase on the stability of virions is evident when the thermal profiles of a single virion type (i.e., B3+4^V^ or C3+4^V^), assembled with homologous vs. heterologous replicase, was comparatively analyzed (Fig. 7F, G). For example, the melting profile of B3+4^V^ virions resembled those of Wt BMV virions, i.e., displaying a significant peak at 49°C and a minor one at 85°C (compare black and blue peaks in Fig. 7D and F, respectively). By contrast, the melting profile of B3+4^V^ virions assembled with the heterologous replicase displayed two prominent peaks: one at 59°C and another at 88°C (Fig. 7F, red peaks). Likewise, the melting profile of C3+4^V^ virions assembled with homologous replicase is distinct from that of heterologous replicase (Fig. 7 E, G).

Next, we tested the sensitivity of B3+4^V^ or C3+4^V^ virions assembled under heterologous replicase to trypsin digestion. Compared to B3+4^V^ virions assembled under the homologous replicase (i.e., p1a+p2a+B3; Fig. 8A, lane 1), a majority of the virions assembled under the heterologous replicase remained resistant to trypsin digestion (i.e., pC1a+pC2a+B3; Fig. 8A, lane 2). By contrast, a majority (∼80%) of C3+4^V^ virions assembled under homologous replicase remained intact (i.e., pC1a+pC2a+C3) while those assembled under heterologous replicase (i.e., p1a+p2a+C3) remained completely resistant (Fig. 8A bottom panel, lane 3) indistinguishable from the undigested control (Fig. 8 top panel, lane 3). EM analysis further substantiated these observations (Fig. 8B).

**FIG 8.**
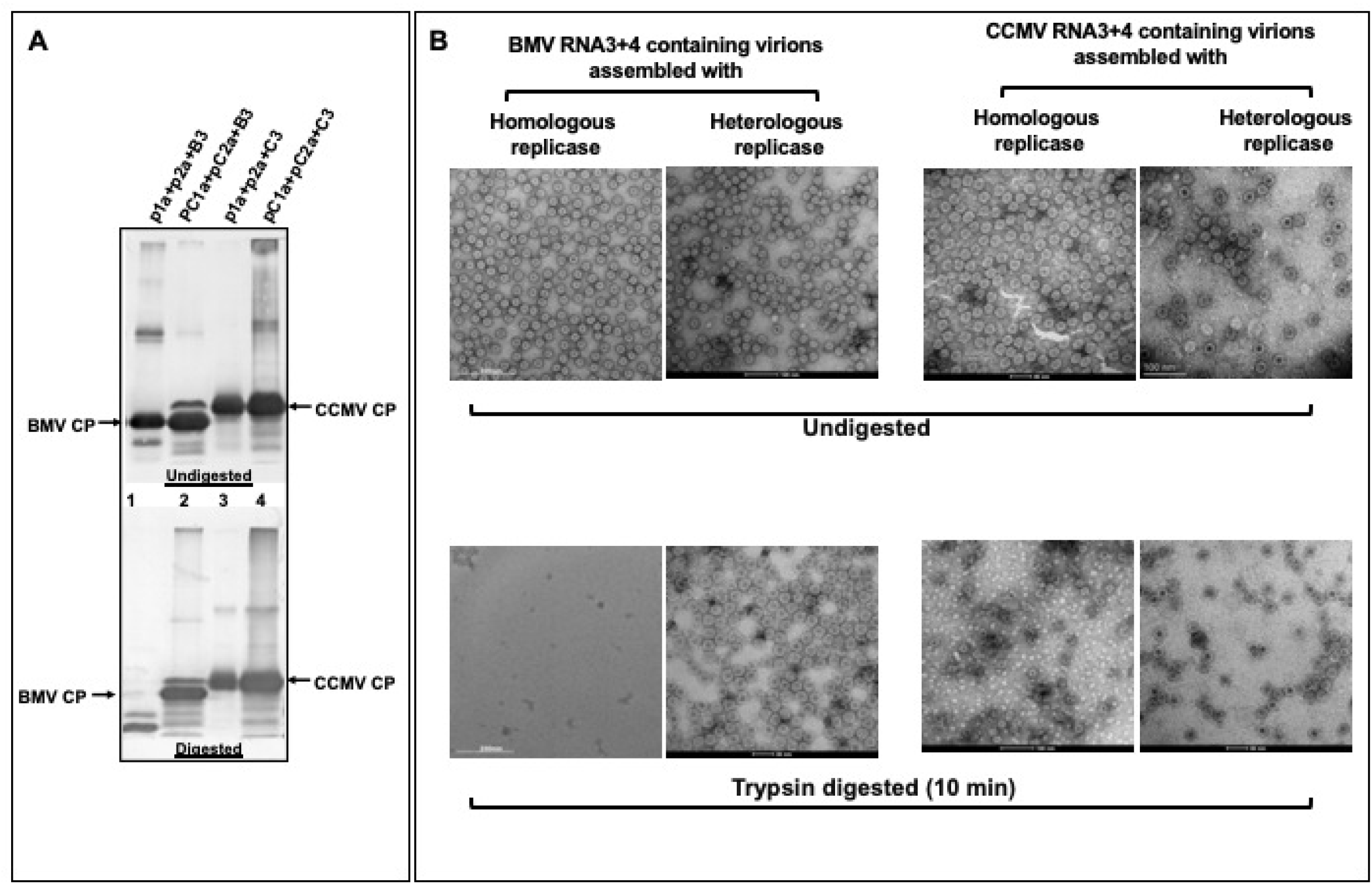
Capsid dynamics of B3+4^V^ and C3+4^V^ virions. (A) Western blot analysis of undigested (control) and trypsin digested B3+4^V^ and C3+4^V^ virions assembled in the presence of either homologous (panel A, lanes 1 and 4) or heterologous replicase (panel A, lanes 2 and 3). (B) Negative-stain electron micrographs showing the integrity of the undigested and trypsin-digested virion preparations are shown in panel A. A size bar is given on each micrograph.

Finally, because of its high sensitivity, MALDI-TOF was used to identify the cleavage products released due to trypsin digestion for B3+4^V^ or C3+4^V^ assembled under homologous and heterologous replicases (Fig. 9). Analysis of the released peptides at early time points (10 min) distinguished B3+4^V^ and C3+4^V^ virions assembled with homologous replicase-CP interaction vs. heterologous replicase-CP interaction. In agreement with the previous report (9), the surface of B3+4^V^ virions assembled with the homologous BMV replicase exposes K65, R103, K111, and K165 residues (Fig. 9A; Table 2). In these virions, although peptides representing N-terminal regions (21 to 26 aa) were detected at the earliest time point (e.g., 10 min) (Fig. 9), their accessibility to protease coincided with other cleavage sites as well (Fig. 9; Table 1). Therefore, it is likely that these N-terminal peptides on virions of B3+4^V^ may not be the first sites of protease cleavage since multiple simultaneous cleavages at other sites expose the otherwise internalized N-terminal region to proteolysis. Detection of peptides encompassing the N-proximal 21 to 26 and 20 to 26 aa region at an early time point (10 min) suggests their externalization in B3+4^V^ (Fig. 9). Alternatively, it is likely that the extensive cleavage of the capsid as early as 10 min by trypsin (Fig. 9) results in degradation of the capsid structure (Fig. 7B), which leads to externalization of the N terminus and its subsequent proteolysis.

**FIG 9.**
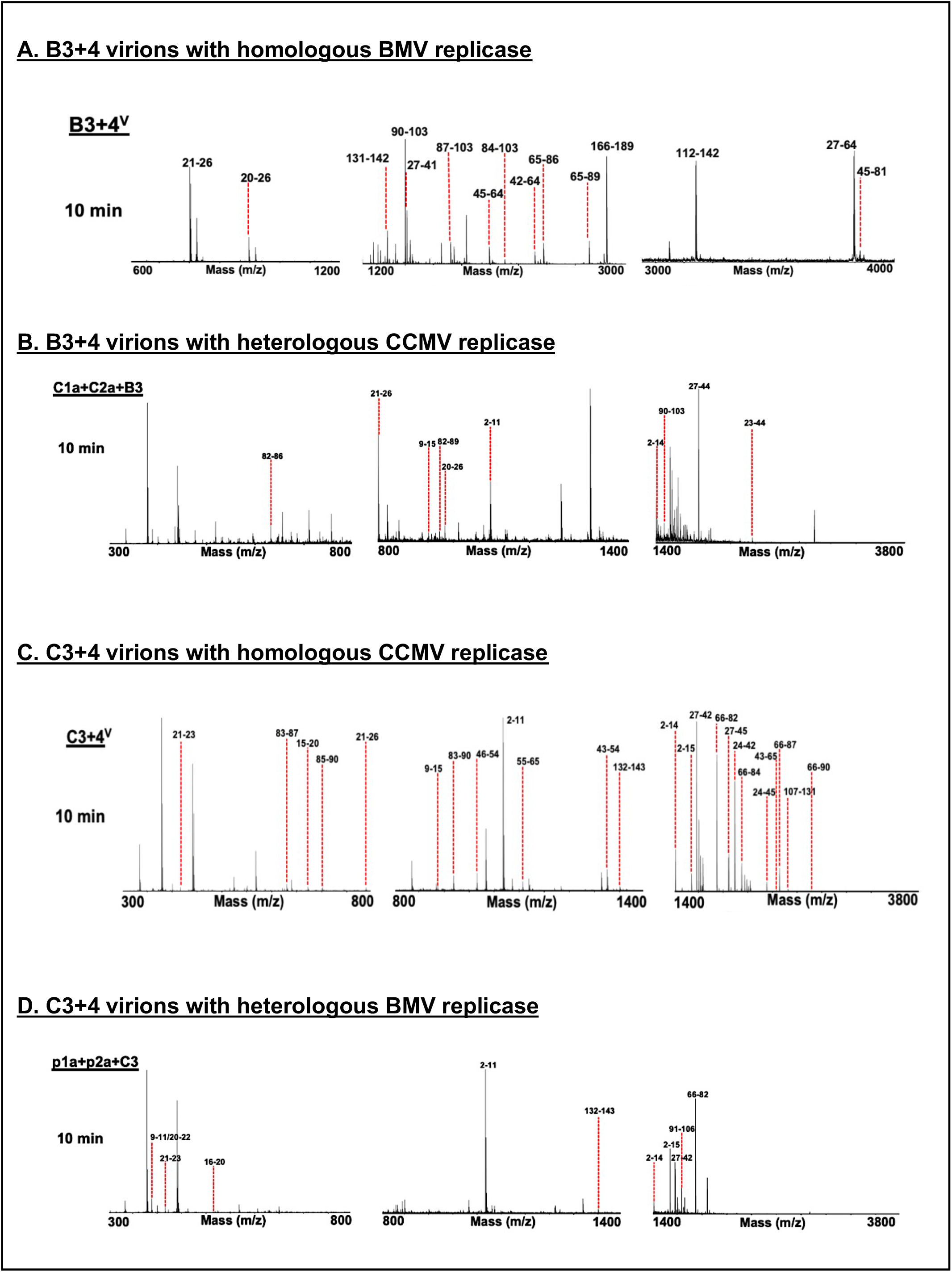
MALDI-TOF analysis of trypsin-digested hybrid virions. (A, B) MALDI-TOF analysis of peptides released from B3+4^V^ virions assembled with homologous and heterologous replicase following digestion with trypsin at 10 min. (C-D) MALDI-TOF analysis of peptides released from C3+4^V^ virions assembled with homologous and heterologous replicase following digestion with trypsin at 10 min. Peaks are labeled with corresponding polypeptide fragments of predicted amino acid residues. Table 2 and 3 summarize masses and identifies the corresponding amino acid residues. The figures indicate representative data from N=2 independent experiments.

**TABLE 2.**
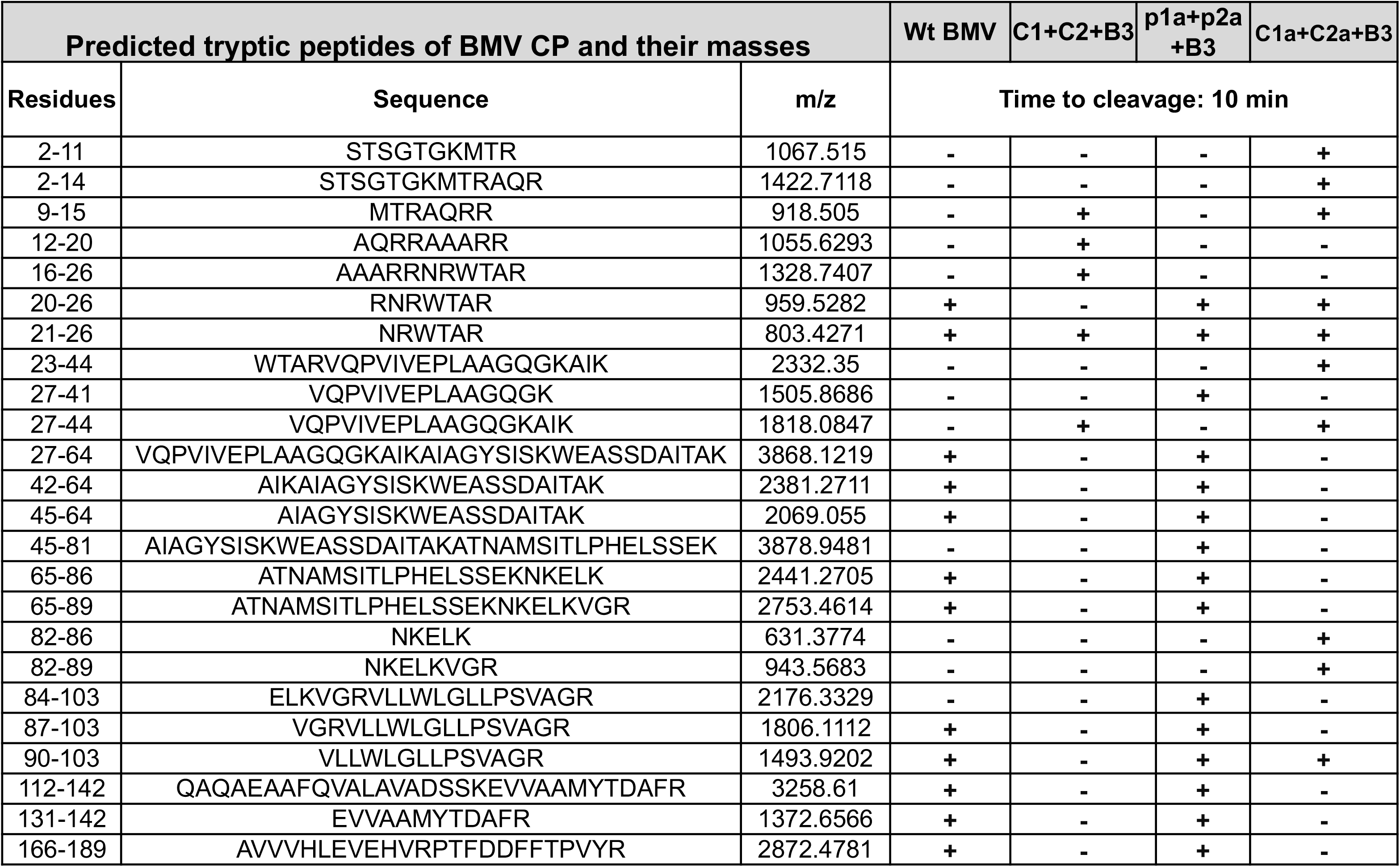
Kinetics of the trypsin cleavage sites located on the Wt BMV and its hybrid virions assembled in the presence of either homologous (Wt BMV and B3+4^V^) or heterologous replicase (C1+ C2+ B3 and C1a+ C2a+ B3). Amino acid peptides fragments recovered for each virion type are indicated with “+.”

In comparison, virions of B3+4^V^ assembled with heterologous replicase (i.e., pC1a+ pC2a+ pB3; Fig. 9B) did not release peptides encompassing previously reported surface residues (K65, K111, and K165) indicating these residues are internalized and not accessible to trypsin cleavage. Also, these virions exposed their N-ARM on the capsid surface, as evidenced by the cleavage products recovered (Fig. 9B; Table 2). However, these residues are not a part of the structure. Therefore, any disruption encompassing them did not impact the overall structural integrity of the virions (Fig. 8B). Table 2 summarizes a detailed profile of the peptide fragments released following trypsin digestion of B3+4^V^ virions assembled under homologs vs. heterologous replicase.

The MALDI profile of trypsin-digested C3+4^V^ assembled under homologous replicase (Fig. 9C) mirror the profile shown in Fig. 5C. Interestingly, the integrity of C3+4^V^ virions assembled under heterologous replicase (Fig. 9D) confirms the release of only negligible amounts of fragments encompassing the N-terminus (Table 3). We presume that a relatively small subset of virions is involved in the release of the low abundance of fragments like 66-82, 91-106, and 131-132.

**TABLE 3.**
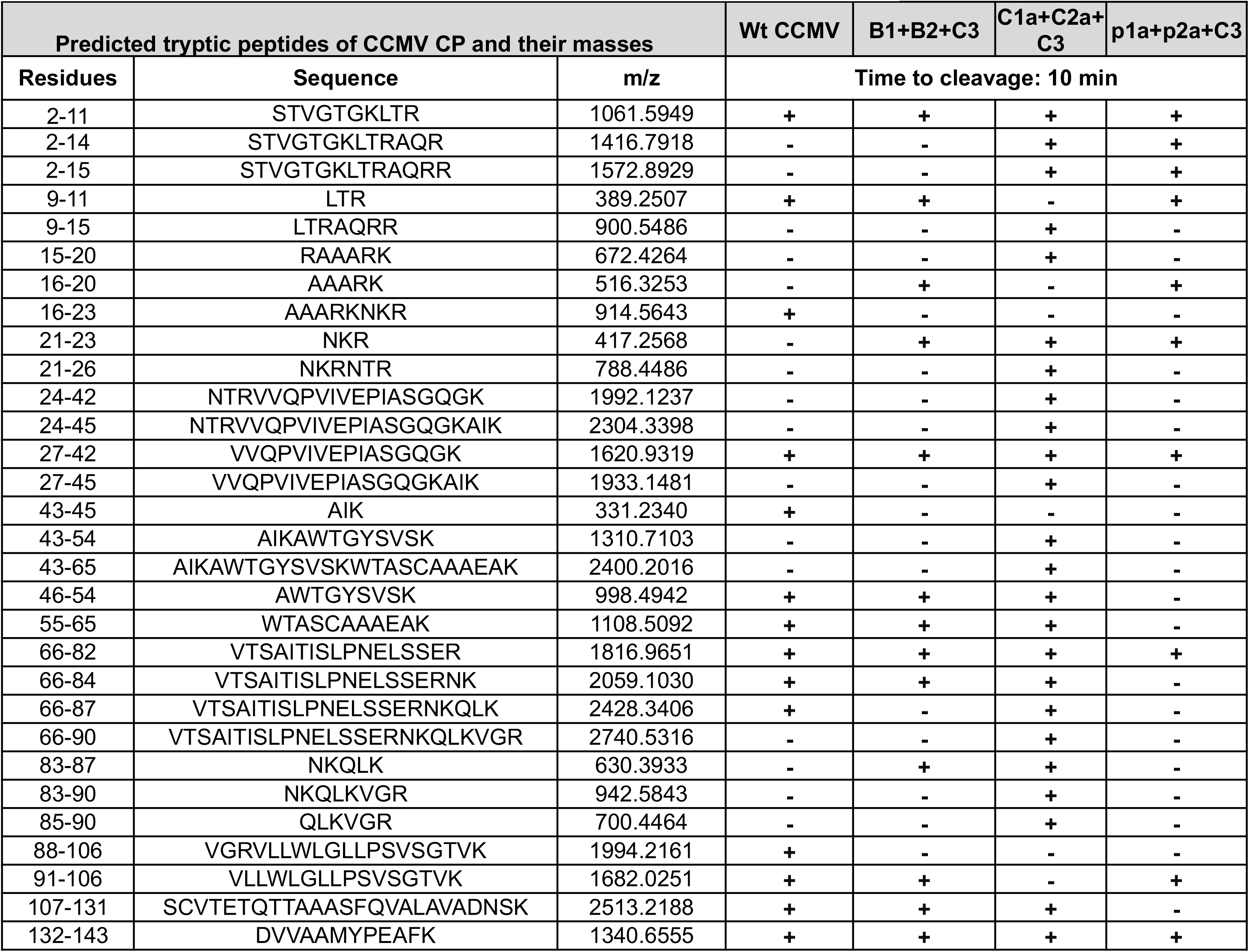
Kinetics of the trypsin cleavage sites located on the Wt CCMV and its hybrid virions assembled in the presence of either homologous (Wt CCMV and C3+4^V^) or heterologous replicase (B1+ B2+ C3 and p1a+ p2a+ C3). Amino acid peptides fragments recovered for each virion type are indicated with “+.”

## Discussion

Despite being similar in the virion structure, genome organization, and replication strategies, CCMV and BMV are biologically distinct: CCMV is adapted to cowpea, a dicot host, while BMV to barley, a monocot host (18). Additionally, both CCMV and BMV also differ in the form of the CP (in conjunction with MP) required for mediating cell-to-cell and long-distance movement (20, 21). Based on this intimacy of CP to host-related activities, we hypothesized that otherwise concealed local and global dynamical conformational changes of the viral capsids are likely to contribute to the host specificity mentioned above. In this study, we extended the agroinfiltration approach (9) to assemble the three virion types of CCMV, C1^V^, C2^V^, and C3+4^V^ (Fig. 1). We analyzed each virions type’s relative stability and capsid dynamics with the application of thermal denaturation analysis followed by peptide-based MALDI-TOF. Subsequently, we evaluated how the type of the host plant and the viral replicase contribute toward modulation of the capsid dynamics. When compared to the previously characterized capsid dynamics of the three virions types of BMV (i.e., B1^V^, B2^V^, and B3+4^V^) to those of CCMV, the results of this study revealed the following: (i) virions of C1^V^ and C2^V^ are indistinguishable in thermal stability and capsid dynamics; (ii) C1^V^ and C2^V^ are dynamically very similar to B1^V^ and B2^V^; (iii) capsid dynamics of C3+4^V^ are different from those of C1^V^ and C2^V^, and B3+4^V^; (iv) the type of host organism had a profound influence on the capsid dynamics of Wt CCMV virions, and finally, (v) virions of B3+4^V^ and C3+^4V^ assembled with heterologous viral replicase noticeably altered the dynamics. Below we discuss the significance of these observations in the context of bromovirus biology.

### Comparative thermal stability and capsid dynamics of the three virion types of CCMV and BMV

Previous biochemical and biophysical studies performed in the last three decades showed that purified virions of Wt CCMV (i.e., a mixture of C1^V^, C2^V^, and C3+4^V^) are morphologically homogeneous and are challenging to fractionate into three distinct virions types (17). Bromoviruses are known to be highly stable at the experimental pH of 4.5 (18). Therefore, at pH 4.5, all three virions types of CCMV did not reveal any difference in their thermal stability (Fig. 3A). However, viral capsid must be flexible to release the packaged genome into the cell to initiate replication followed by infection. Consequently, at pH 7, a physiological pH of the plant cell, bromoviruses have been shown to dissociate into dimers (19). DSF analysis at pH 7.2 revealed variations in the thermal stability of three virion types of CCMV (Fig. 3 A, B). Therefore, the varied instability of C1^V^/C2^V^ vs. C+4^V^ (Fig. 3, A and B) in conjunction with altered capsid dynamics are implicated in the infection process (see below).

#### Virion types of C1^V^/C2^V^ and B1^V^/B2^V^ are dynamically similar

Concerning trypsin sensitivity, virion types 1 and 2 (i.e., C1^V^/C2^V^ and B1^V^/B2^V^) did not reveal any clear distinction. For example, as observed with B1^V^ and B2^V^ (9), trypsin digestion did not affect the integrity of C1^V^ and C2^V^ virions (Fig. 4A, B) despite the release of 2-15 amino acids encompassing the N-terminal proximal region (Fig. 5 F-G; Table 1). It is not surprising since, in CCMV and BMV, the first N-proximal 25 amino acid region is not visible in the crystal structure and is internalized by interacting with RNA during encapsidation (20, 21). Therefore, trypsin cleavage followed by MALDI-TOF analysis suggested that the capsid surface transiently exposes the N-proximal region, and its release did not affect the integrity of the C1^V^ and C2^V^. Therefore, since the genomic RNAs encapsidated by C1^V^ and C2^V^ encode replicase proteins, the transient exposure of the N-proximal region of C1^V^ and C2^V^ is linked to the co-translational disassembly mechanism, as previously hypothesized for B1^V^ and B2^V^ (9).

#### (i) Virion types of C3+4^V^ and B3+4^V^ are dynamically distinct

A close comparison of trypsin sensitivity profiles of virions of C3+4^V^ and B3+4^V^ displayed a scenario contrasting to that of C1^V^/C2^V^ and B1^V^/B2^V^. The partial and complete disruption of virion integrity for C3+4^V^ (Fig. 4A, B) and B3+4^V^ (9) support these observations. In bromoviruses, virion assembly requires dimerization of the N- and C-terminal CP subunits, and the C-terminal region plays a critical role in this process (16). Consequently, exposure of the C-terminal region to trypsin as early as 10 min is sufficient to abolish the virion integrity. However, a significant difference between the C3+4^V^ vs. B3+4^V^ is the relative accessibility of the C-terminus for trypsin digestion. For example, since a significant percentage of virions remain intact (Fig. 4B) suggests that in C3+4^V^, only a certain percentage of virions expose the C-proximal region for trypsin digestion. Therefore, it is reasonable to assume the segregation of C3+4^V^ virions into two distinct populations. A major population is resistant to trypsin (due to inaccessibility of the critical C-terminal peptides involved in the CP dimerization), and a minor population is susceptible to trypsin (due to transient exposure of the C-terminal peptides). By contrast, following trypsin digestion, the integrity of the entire population of B3+4^V^ virions was completely disrupted (9).

#### Then, how do these two contrasting scenarios of C3+4^V^ vs. B3+4^V^ contribute to the life cycles of CCMV and BMV?

We offer the following explanation. In bromoviruses, gene products encoded by RNA3 play essential roles in replication and host adaptability (22). In CCMV and BMV, CP mRNA production depends on the replication of the genomic RNA3 (4, 23). Following translation, CP stimulates the asymmetric synthesis of the plus-strand over the minus strand (24), and its interaction with replicase dictates packaging specificity (25). Therefore, based on the capsid dynamics of B3+4^V^, we previously hypothesized that the complete release of the genomic RNA3 is obligatory for its replication (9). However, this study revealed that, unlike B3+4^V^, most C3+4^V^ virions are stable. Taken together, we speculate that virions of C3+4^V^ and B3+4^V^ evolved to modulate their capsid dynamics during *in vivo* assembly to adapt to their respective natural hosts and the CP form required for the cell-to-cell movement.

### Modulation of the capsid dynamics by the host organism

Studies analyzing the capsid dynamics of RNA viruses pathogenic to animals and plants offer new insight into how the capsid dynamics fluctuations would control the overall infectivity and pathogenesis (26). Results of this study analyzing the capsid dynamics of CCMV virions obtained from cowpea and *N. benthamiana* host plants (Fig. 6 A-D) accentuates the significance of the viral capsid’s flexibility in establishing a successful infection. Viral infection in a given host plant involves two active processes. These are cell-to-cell and long-distance movement. Following entry and initial replication in primary cells, all plant viruses must spread to the neighboring cells (i.e., cell-to-cell movement) via plasmodesmata with the help of a dedicated and virus-encoded MP (27). However, in BMV and CCMV, both MP and CP are obligatory for cell-to-cell spread (4). A majority of plant viruses, including bromoviruses, manifesting the long-distance spread require assembled virions. These observations suggest that an optimal interaction between the host machinery and the virion surface configuration (arrangement of amino acids) is necessary to mediate the long-distance spread. A change in the virion surface configuration due to a mutation may lead to altered spread and pathogenesis. For example, BMV induces chlorotic local lesions on inoculated leaves followed by systemic mottling symptom phenotype in *Chenopodium quinoa* (10). A small deletion in the N-proximal region significantly altered the lesion phenotype and blocked the systemic spread (10). Given the variation in the cell types of cowpea and *N. benthamiana*, considerable flexibility in capsid dynamics is envisioned for CCMV. This required flexibility perhaps explains why the capsid dynamics of CCMV are distinct in cowpea and *N. benthamiana* plants (Fig. 6).

### Modulation of capsid dynamics by the viral replicase

In addition to catalyzing replication, viral replicase is intimately involved in regulating other biological processes such as pathogenesis and genome packaging. In BMV and CCMV, the assembly of biologically viable hybrids involves exchanging genomic RNA3 (i.e., C1+C2+B3 and B1+B2+C3), but not either genomic RNA 1 or RNA 2 (e.g., C1+B2+C3; B1+C2+C3) (15). Furthermore, the interaction of viral replicase with the CP alters the RNA packaging specificity (25). Results of this study have identified another critical role played by the viral replicase, i.e., modulation of capsid dynamics reflecting the stability. The effect of viral replicase on the stability and capsid dynamics became apparent since virion types of BMV and CCMV assembled under the direction of homologous and heterologous replicase displayed contrasting profiles (Figs. 7-9; Tables 2 and 3).

Based on the MALDI-TOF data (Tables 1-3), how the dynamics of B3+4^V^ and C3+4^V^ virions are modulated by the homologous and heterologous replicase is schematically shown in Fig.10. Linear CP of BMV and CCMV respectively expose approximately 24 and 30 trypsin cleavage sites [Fig. 10, Panels A, B (i)]. In the case of BMV, the interaction of the CP subunits with the homologous replicase results in the assembly of B3+4^V^ virions with a confirmation presenting 15 of the 24 trypsin cleavage sites on the surface [Fig. 10, Panel A (ii)]. This conformation rendered B3+4^V^ virions highly susceptible to trypsin and affected the structural integrity (Fig. 8A, lane 1). By contrast, the assembly of B3+4^V^ virions under heterologous replicase altered the surface conformation presenting 10 of the 24 trypsin cleavage sites on the surface [Fig. 10, Panel A (iii)]. This configuration rendered B3+4^V^ virions resistant to trypsin (Fig. 8, lane 2).

**FIG 10.**
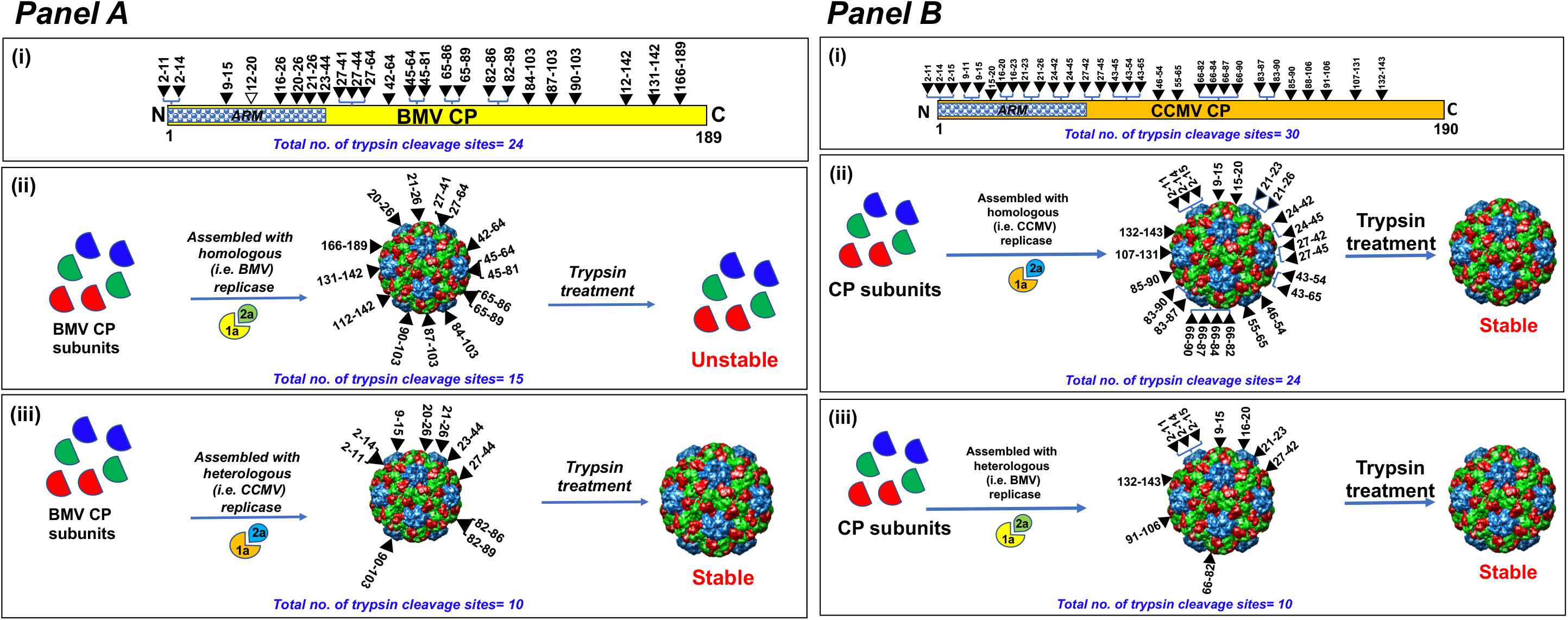
A schematic model showing the alteration of trypsin cleavage on the surface of B3+4^V^ and C3+4^V^ assembled under the homologous vs. heterologous replicase (see text for details).

With respect to CCMV, the interaction of the CCMV CP subunits with the homologous replicase results in the assembly of C3+4^V^ virions with a confirmation presenting approximately 24 of the 30 trypsin cleavage sites on the surface [Fig. 10, Panel B (ii)]. This confirmation rendered ∼20% of C3+4^V^ virions susceptible to trypsin while the remaining 80% remained structurally intact (Fig. 8). Whereas C3+4^V^ virions assembled under the heterologous replicase positioned 10 of the 30 trypsin cleavage sites and rendered 100% virions resistance to trypsin [Fig. 10, Panel B (iii)]. The contrasting trypsin sensitivity of B3+4^V^ and C3+4^V^ virions assembled under homologous vs. heterologous replicase (Figs. 8-10) provides compelling evidence for the intimate involvement of the viral replicase in modulating the capsid dynamics. Apparently, interaction with the viral replicase positions the biologically relevant peptides on the capsid surface for optimal interaction with the host for successful onset of the infection. These observations explain why neither hybrid virus (i.e., C1+C2+B3 or B1+B2+C3) systemically infected the natural host of CCMV and BMV (15).

In conclusion, this study accentuates previously unrecognized structural differences among the three virions of CCMV and BMV and their importance in the general biology of the two viruses. The previously determined cryo-structure of Wt CCMV represents the average of three virions (20). Therefore, the strategy used in this study (Fig. 1) would be ideal for comparatively assessing the structure of independent virion types. Such studies are likely to shed more light on the structural and conformational heterogeneity of the three virions types and their role in viral biology and pathogenesis.

## Materials and Methods

### Agro-plasmids used in this study

The construction and characteristic properties of agro-plasmids pC1, pC2, and pC3 (Fig. 1, Panel IA), engineered to express biologically active full-length genomic RNAs of CCMV following agroinfiltration into plants, are described previously (9). Likewise, agro-plasmids pC1a, pC2a, and pCCP (Fig. 1, Panel IB) are engineered respectively to transiently express replicase protein1a (p1a), replicase protein 2a (p2a) and CP are as described previously (9). Agro constructs for expressing the full-length BMV genomic RNAs (Fig. 1, Panel IC) and for transiently expressing the replicase, and CP (Fig. 1, Panel ID) are previously described.

### Virion purification, electron microscopy (EM), virion RNA electrophoresis, and Western blot analysis

The procedure used to purify CCMV and BMV virions from 4-6 days post-infiltration (dpi) agroinfiltrated leaves or mechanically inoculated cowpea (for CCMV) and *N. benthamiana* (both CCMV and BMV) was as described by Rao et al. (28). For negative-stain EM analysis, gradient purified virions were spread on glow-discharged grids followed by negative-staining with 2% uranyl acetate prior to examination with a Technai 12 transmission electron microscope operated at 120KeV (Center for Advanced Microscopy and Microanalysis facility at UC-Riverside) and images were recorded digitally. For electrophoretic mobility analysis, purified virions were loaded in a 1% agarose gel prepared and electrophoresed in virus suspension buffer at 7v/cm for 2-2.5 hr at 4^0^C. Western blot analysis of un-digested and trypsin-digested virion samples using the desired anti-CP (either anti-CCMV CP or a mixture of both anti-BMV CP and anti-CCMV CP) was performed as described previously (29).

### Differential scanning fluorimetry (DSF)

DSF was performed essentially as described previously (30). Desired virus and control (lysozyme) samples were re-suspended either in nano pure sterile water (pH 7.1) or in virus suspension buffer (50 mM sodium acetate, 8 mM magnesium acetate, pH 4.5) or in 100 mM Phosphate buffer (pH 7.2). The experiment was performed three times independently using three replicates for each sample. DSF data were analyzed as described by Rayaprolu *et al*. (30) and plotted using GraphPad Prism version 9.3.1 for macOS, GraphPad Software, San Diego, California USA, www.graphpad.com.

### MALDI-TOF

For MALDI-TOF analysis, a desired purified virion preparation was diluted to 1 mg/ml in 25 mM Tris-HCl, 1 mM EDTA buffer. A 30 μl sample (i.e., 30 μg of the virus) was digested with 1:100 (w/w) ratios of trypsin (Pierce Trypsin Protease mass spectrometry grade; Thermo Fisher Scientific)/virus for various time points at 25°C (31, 32). A 20 μl of water was added to 10 μl of the digested sample, and approximately 0.5 μl of each digest or undigested (control) sample was analyzed on AB Sciex TOF/TOF 5800 MALDI MS with a-cyano-4-hydroxycinnamic acid matrix. The instrument was calibrated with standards, and the test samples were analyzed with external calibration. The accuracy is about +/- 0.05 dalton. The MALDI-TOF data were analyzed with AB Sciex Data Explorer, and baseline was corrected with peak width 32, flexibility 0.5 and degree 0.1. For noise removal, the standard deviation was set at 2. To determine the peak detection criteria, % centroid was taken as 50, signal/noise (S/N) threshold was 3, and the noise window width m/z was 250. The threshold after S/N recalculation was set at 20. The peptide fragments were assigned based on the UCSF Protein Prospector’s MS-Digest function (33).

## ACKNOWLEDGMENTS

We thank Matthew Dickson at the Center for Advanced Microscopy and Microanalysis facility at UC-Riverside (UCR) for helping with imaging the virions, Jie Zhou at Analytical Chemistry Instrumentation Facility at UCR for helping with MALDI-TOF analysis, and Institute for Integrative Genome Biology at UCR for letting us use the facility for DSF experiment.

This research was supported by grants from the UC AES/RSAP (19900) and Academic Senate.

